# New genetic tools in *Finegoldia magna* identify a conserved adhesin required for the formation of stress-tolerant aggregates

**DOI:** 10.64898/2026.09.18.752566

**Authors:** Susannah Lawhorn, Cassandra J. Speyer, Angelie Arcos Chavelas, Sierra R. Scamfer, Francia Lopez Palomera, T. Jarrod Smith, Alison Coluccio, Melanie A. Spero

## Abstract

The Gram-positive obligate anaerobe *Finegoldia magna* is a member of the healthy human microbiota, but also acts as an opportunistic pathogen to cause persistent, biofilm-associated infections. Despite its prevalence on the human host, little is known about *F. magna* biology or the mechanisms underlying its clinically relevant phenotypes, partly due to the lack of genetic tools. We address this gap by establishing genetic approaches for investigating gene function in *F. magna*, which we apply to identify genetic determinants of aggregate biofilm formation. We found that *F. magna* isolates are naturally competent, allowing for targeted chromosomal integration of linear DNA constructs via homologous recombination. Transformation frequency varied substantially among strains and was also affected by factors such as homologous flank length, DNA concentration, and incubation method. To identify genes that mediate aggregate biofilm formation, we used experimental evolution to select for *F. magna* mutants that had lost the ability to aggregate. This approach identified a conserved locus encoding a putative adhesin that we named FafA (*<u>F</u>inegoldia* <u>a</u>dhesion factor <u>A</u>). Next, we applied targeted mutagenesis tools to show that deletion of *fafA* markedly reduced autoaggregation but does not impair other modes of biofilm formation, including surface attachment or aggregation during agitation. Finally, we demonstrate that FafA-mediated aggregation protects *F. magna* from antibiotic and oxidative stress. Together, these findings establish a genetic framework for mechanistic studies in *F. magna* and identify FafA as a conserved adhesin that promotes aggregation and stress tolerance in this anaerobic pathobiont.

**Importance:** Despite growing recognition of *Finegoldia magna* as both an important member of the human microbiota and opportunistic pathogen, mechanistic studies of this organism have been limited by the lack of genetic tools. Here, we establish a system for targeted mutagenesis that leverages this bacterium’s natural ability to take up DNA from its environment. We combine these new tools with a forward-genetics approach to identify the genetic basis of aggregation, a form of biofilm growth underlying recalcitrant infections. These approaches identified the putative adhesin FafA, which mediates a specific biofilm lifestyle that promotes tolerance to antibiotic and oxidative stress. These findings provide a genetic framework for studying *F. magna* biology and offer new insight into the mechanisms promoting biofilm-associated survival.

## Introduction

*Finegoldia magna* is a Gram-positive obligate anaerobe implicated in human health and disease. As a member of the normal human microbiota, *F. magna* colonizes the skin, oral cavity, and gastrointestinal and urogenital tracts (1). Yet it also acts as an opportunistic pathogen, causing a wide range of both acute and chronic infections (2). In the clinic, *F. magna* is most frequently isolated from soft tissue abscesses, wound infections, and bone and joint infections (3). Its capacity to cause both acute and persistent infections is thought to be facilitated by an array of virulence factors, including proteins that degrade host tissues, modulate or evade immune responses, and mediate adhesion to host surfaces (4, 5).

*F. magna* is increasingly recognized for its role in persistent infections, which are often characterized by biofilm formation. In biofilms, bacteria grow adhered to one another as multicellular aggregates or attached to surfaces such as implanted medical devices (6). Biofilm growth is a hallmark of difficult-to-treat infections, since biofilms are resistant to host immune defenses and highly tolerant to antibiotic treatment (7). *F. magna* is one of the most common anaerobes isolated from prosthetic joint infections, often as the sole infectious agent (8–11), and it also colonizes other implanted medical devices such as stents (12, 13) and prosthetic heart valves (14, 15). Biofilm growth is also common in chronic wound infections, where aggregate biofilms are associated with delayed wound healing (16–18). *F. magna* is among the most prevalent genera colonizing chronic wounds (19), where increased anaerobe abundance in wounds is correlated with worse patient outcomes, including more severe infection (20), longer-lasting wounds (21–23), and amputation (24). Further supporting a role for biofilm-mediated persistence, *F. magna* failed to persist in a mouse wound model when introduced as planktonic cells, but remained abundant when introduced as part of a polymicrobial biofilm that impaired wound healing (25). In laboratory conditions, many *F. magna* strains readily aggregate in culture, and prior work identified a surface protein predicted to promote *F. magna* adherence to ex vivo human skin samples (26). Taken together, *F. magna* exhibits multiple clinically relevant modes of biofilm growth, yet the genetic determinants underlying its biofilm formation and persistence remain largely undefined.

Despite its clinical significance, *F. magna* biology has been underexplored for several reasons. Clinical laboratories do not routinely perform anaerobic culturing, contributing to the underdiagnosis of anaerobic pathogens and ultimately limiting recognition of their clinical significance (27). Even deliberate efforts to culture anaerobes often fail to recover them, as they require specialized equipment and many species are slow-growing and fastidious (3). As a result, culture-based methods frequently underrepresent anaerobic taxa in clinical samples; however, sequencing-based approaches often reveal anaerobes to be abundant in samples where they were not detected by culture (22, 28, 29). Additionally, obligate anaerobes are common members of polymicrobial infections (2). In such instances, anaerobes are often overlooked in favor of community members that are readily culturable and deemed more clinically significant (e.g. facultative anaerobes like *Staphylococcus aureus* or *Pseudomonas aeruginosa*). Finally, a lack of research tools for studying anaerobes like *F. magna* has hindered mechanistic studies of the processes that underpin its colonization, biofilm lifestyles, and persistence.

In this work, we establish genetic tools for *F. magna* and apply them to investigate biofilm formation. These tools leverage natural competence in *F. magna* and were adapted from recent work in the related Gram-positive anaerobe, *Parvimonas micra* (30). After optimizing transformation efficiency across different *F. magna* isolates, we used this approach to generate targeted gene deletion mutants. We further used experimental evolution to identify a novel adhesin that we named FafA (*<u>F</u>. magna* <u>a</u>dhesion factor <u>A</u>), which mediates static autoaggregation but not other modes of biofilm growth, indicating that *F. magna* employs multiple genetic systems to control distinct biofilm programs. We further show that FafA-dependent aggregation promotes *F. magna* tolerance to antibiotic and oxidative stress, as is characteristic of biofilms. FafA is widely conserved across the *F. magna* lineage and also present in other Gram-positive anaerobes. Together, our work establishes genetic tools for future mechanistic studies of *F. magna* biology, and highlights the importance of the biofilm lifestyle for this anaerobic pathobiont.

## Results

### *F. magna* isolates are naturally competent for transformation

To date, the only report of targeted genetics in *F. magna* is the construction of a gene disruption strain that was generated by introducing a suicide vector via electroporation (31). To our knowledge, this method has not since been used by others, and we were unsuccessful adapting this approach for *F. magna* reference strain ATCC 29328. Thus, we sought another method for targeted mutagenesis in *F. magna*.

Higashi et al. (30, 32) recently developed genetic tools for *Parvimonas micra* by exploiting its natural competence. Due to the relatedness of these organisms, we investigated whether methods employed in *P. micra* could be adapted for *F. magna*. To test for natural competence, we PCR-amplified and assembled a DNA construct comprising an erythromycin resistance cassette (*ermB*) flanked by 2 kb fragments that are homologous to sites downstream of the *F. magna tuf* gene (Fig. 1A), similar to Higashi et al. *F. magna* cultures were mixed with 1 µg of the DNA construct, spotted onto THB-Tween blood agar, and incubated overnight. After incubation, cells were scraped, resuspended in PBS, and plated onto THB-Tween blood agar with or without erythromycin to determine transformation frequency. By performing transformations in four *F. magna* isolates, we determined that all strains exhibit natural competence, but that transformation frequency varied greatly between strains, ranging almost 4 orders of magnitude (Fig. 1B). Next, we conducted whole genome sequencing on transformed isolates from each strain, which confirmed that *tuf-ermB* knock-in mutants were successfully generated in each strain background. Interestingly, sequencing results also revealed that the transformation process caused *F. magna* ATCC 29328 to lose its native 190 kb plasmid, pPEP1 (33). *F. magna* ATCC 29328 is the only strain in these studies that maintains a native plasmid, and we consistently observed plasmid loss when constructing other mutations in this background (described below).

**Figure 1.**
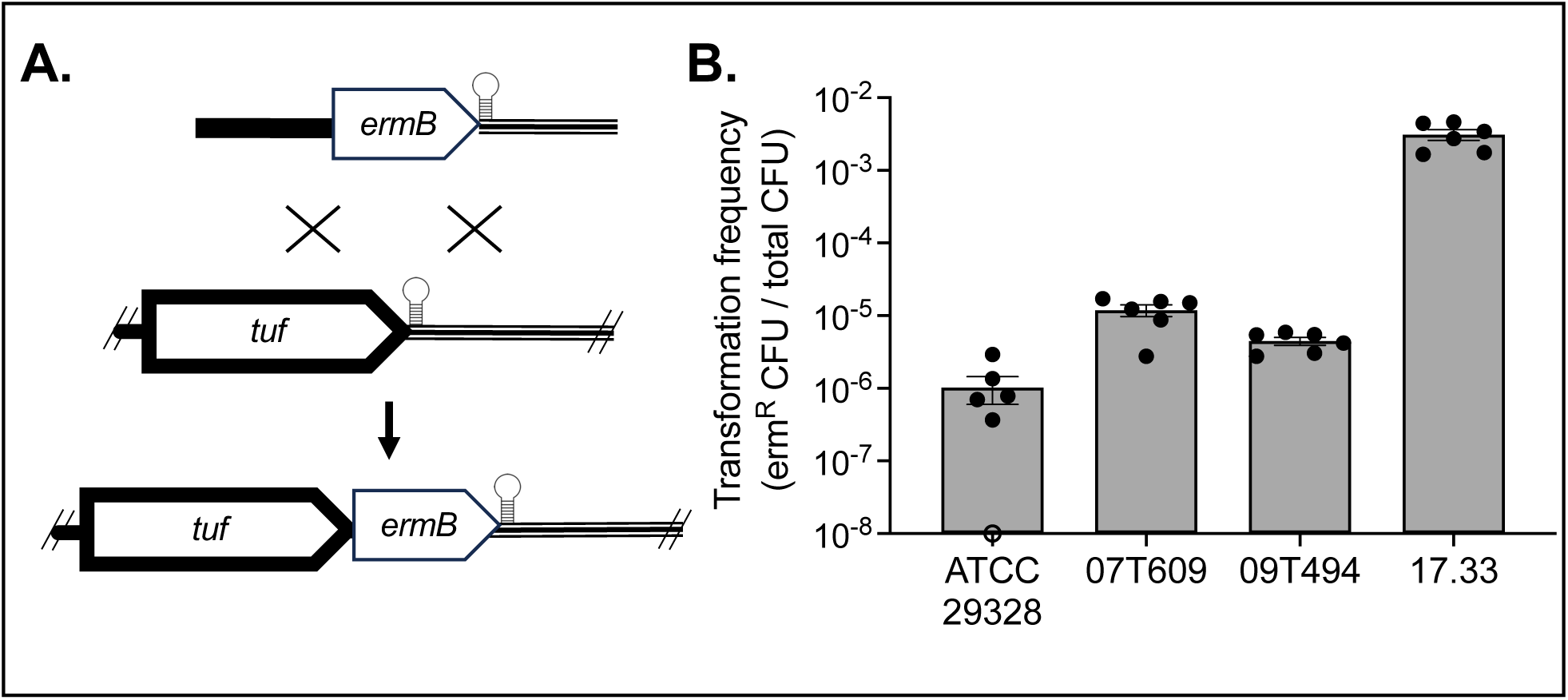
Transformation frequency varies across *F. magna* isolates. **A.** Mutagenesis construct (top) containing an erythromycin resistance cassette (*ermB*) was inserted into the *F. magna* genome downstream of the *tuf* ORF but upstream of the *tuf* terminator (loop) via homologous recombination (middle), incorporating *ermB* into the genome at that locus (bottom). **B.** Transformation frequency for four *F. magna* isolates was determined as the ratio of erythromycin-resistant CFU to total CFU. Shown are two technical replicates from three independent experiments with the mean ± standard error of the mean (SEM). Open circles on the x-axis indicate transformation frequency below the detection limit of the assay.

### Optimal transformation conditions vary between *F. magna* isolates

We next aimed to optimize transformation in *F. magna* by identifying variables that influence transformation frequency. For these experiments, we focused on two strains of *F. magna*, ATCC 29328 and 17.33, representing isolates with the lowest and highest transformation frequencies, respectively (Fig. 1B). First, we tested the impact of homologous flanking region size, using mutagenesis constructs with 500 – 2500 bp homologous flanks (Fig. 2A). Increasing the size of the homologous DNA fragment increased transformation frequency in both ATCC 29328 and 17.33, with highest frequencies observed for constructs with 2000 – 2500 bp flanks (Fig. 2B). However, transformation frequencies in ATCC 29328 remained orders of magnitude lower than the highest frequencies in 17.33.

**Figure 2.**
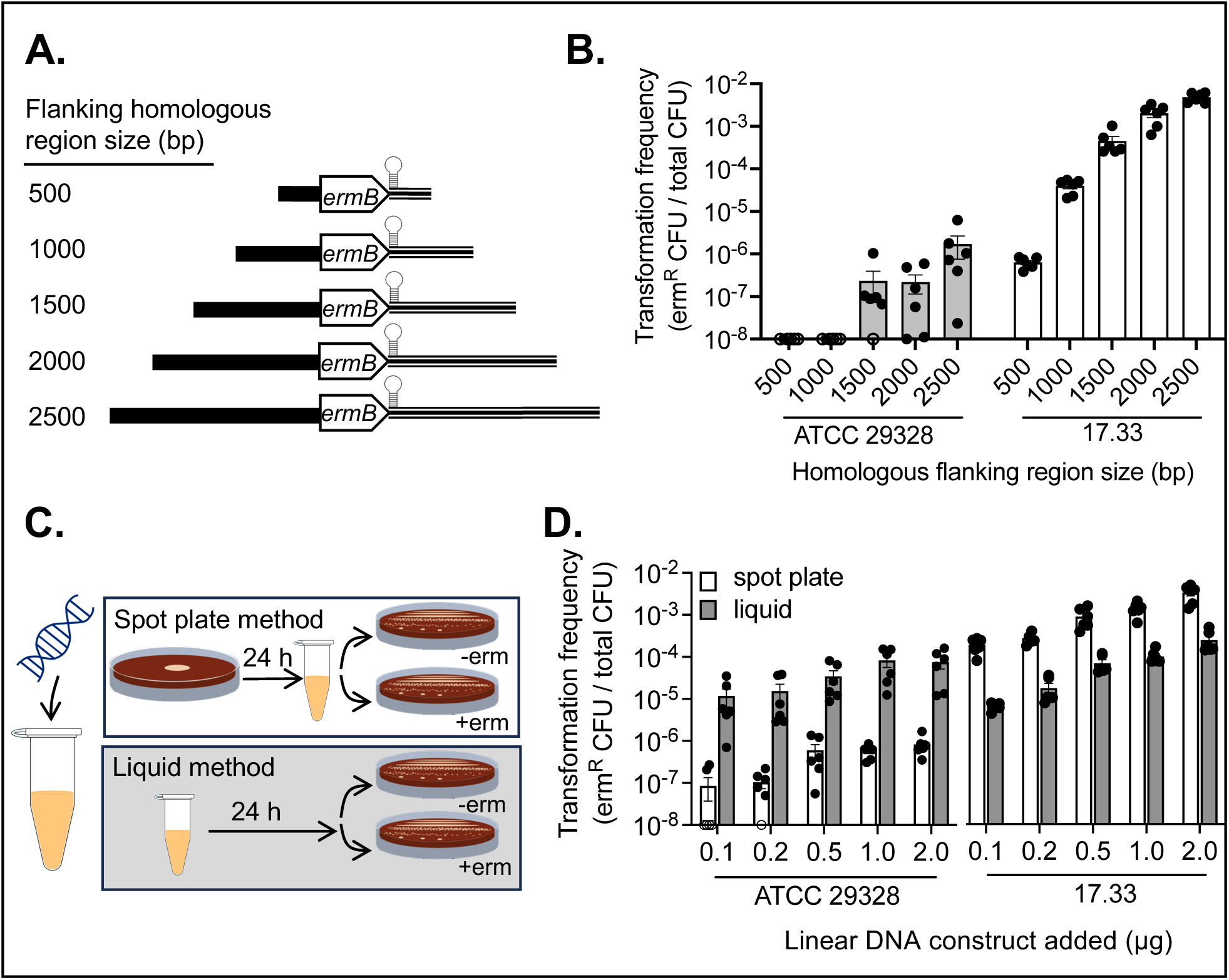
Optimal transformation conditions differ between *F. magna* isolates. **A.** Mutagenesis constructs, inserting an erythromycin resistance cassette (*ermB*) downstream of the *tuf* ORF but upstream of the *tuf* terminator (loop), were generated with homologous flanking regions of various sizes. **B.** Strains ATCC 29328 and 17.33 were transformed with these constructs and transformation frequency was determined as the ratio of erythromycin-resistant CFU to total CFU. **C.** During transformation, *F. magna* cells were incubated with DNA constructs for 24 hours either on a plate or in liquid, after which cells were plated with or without erythromycin to quantify transformation frequency. **D.** ATCC 29328 and 17.33 were transformed using each incubation method across a range of DNA construct concentrations and transformation frequency was determined. Shown in B. and D. are two technical replicates from three independent experiments with the mean ± SEM, where open circles on the x-axis indicate transformation efficiency below the detection limit of the assay.

To improve transformation frequencies, particularly in ATCC 29328, we next tested the effect of DNA concentration and incubation method. In previous experiments, cells were mixed with 1 µg of DNA and spotted onto a THB-Tween blood agar plate for overnight incubation, after which cells were scraped and plated onto media with or without erythromycin; we refer to this approach as the spot plate method. For these experiments, we incubated cells with DNA concentrations ranging from 0.1 – 2.0 µg under one of two incubation conditions: via the previously used spot plate method or via a liquid incubation method where cell cultures and DNA were incubated overnight in liquid THB-Tween (Fig. 2C). Increasing DNA concentration had modest effects in increasing transformation frequency for both strains (Fig. 2D). Interestingly, the two isolates exhibited different responses to the incubation methods. For 17.33, the spot plate method consistently outperformed the liquid incubation method, whereas the liquid incubation method yielded higher transformation frequencies for ATCC 29328 (Fig. 2D). Critically, the liquid incubation method increased transformation frequencies by ∼100-fold for ATCC 29328. Together, these results indicate that incubation method is a consequential factor for transformation outcomes in *F. magna* and that the optimal protocol differs between strains.

### Identification of a putative adhesin involved in aggregate biofilm formation

Having developed an approach for targeted mutagenesis in *F. magna*, we next sought to use these tools to investigate its physiology. Given this organism’s propensity for biofilm growth and the clinical significance of biofilms, we aimed to uncover genetic determinants of biofilm formation in this opportunistic pathogen. When *F. magna* is cultured statically in THB-Tween, it will autoaggregate: gentle agitation of an overnight culture causes cells to form a large aggregate that settles at the bottom of the culture well (Fig. 3A). Here, we used experimental evolution to identify genes required for aggregation, an approach that has been successful in other organisms (34–36). We repeatedly passaged the planktonic fraction of an *F. magna* ATCC 29328 culture, avoiding the settled aggregate, to select for mutants that were unable to aggregate. Across 10 lineages, all cultures became increasingly planktonic over time, with cultures largely losing their ability to grow as an aggregate by passage 9 (Fig. 3B). We streak purified isolates from five lineages at passage 10 and conducted whole genome sequencing. All isolates harbored mutations at a locus comprised of the uncharacterized genes, FMG_0048 and FMG_0049 (Fig. 3C, Table S1).

**Figure 3.**
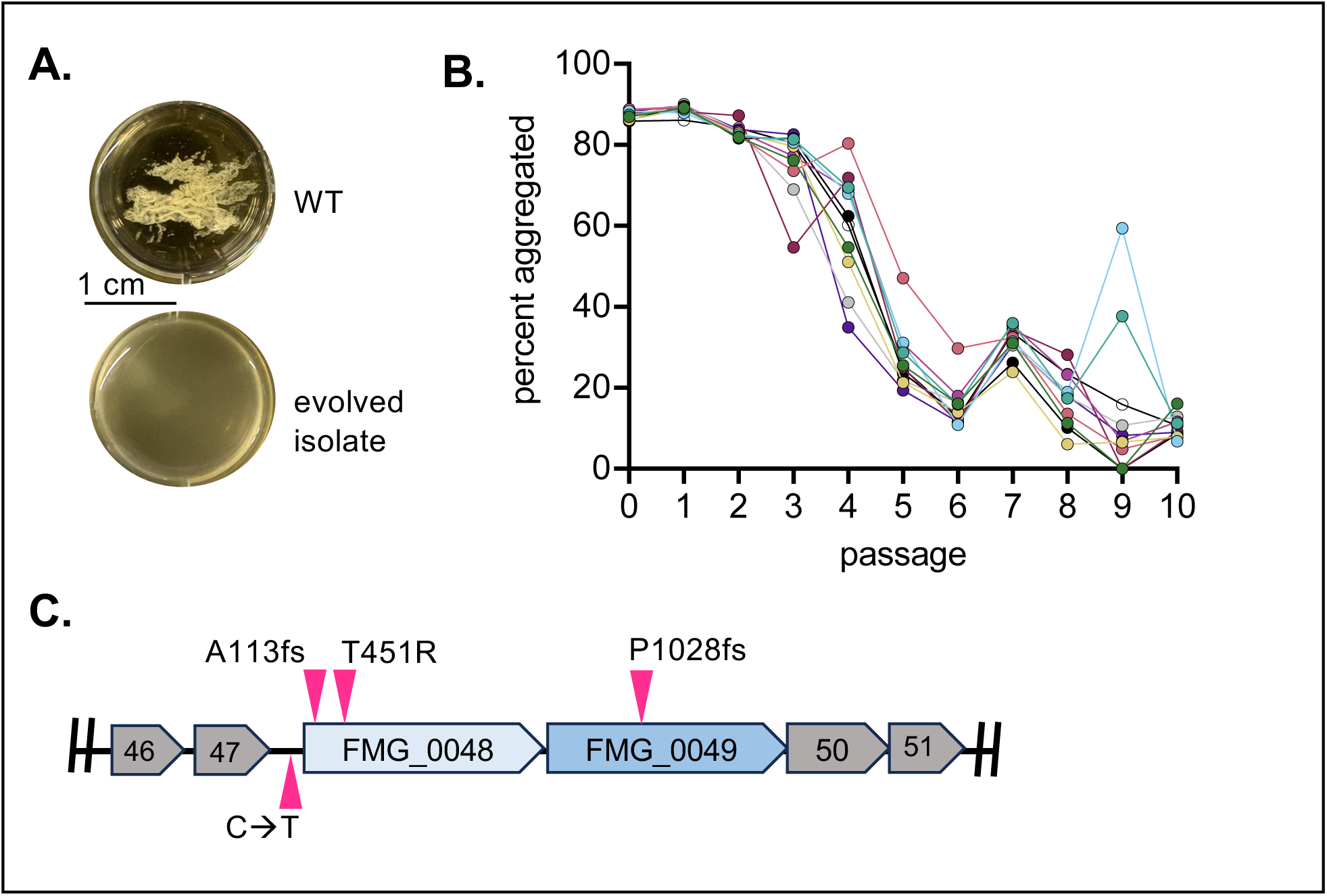
Experimental evolution identifies a locus involved in autoaggregation. **A.** Statically grown overnight cultures of WT *F. magna* ATCC 29328 (top) autoaggregate after gentle agitation (imaged in a 12-well plate), whereas experimentally evolved isolates grow planktonically (bottom). **B.** Experimental evolution was conducted in 10 independent lineages, serially passaging only the planktonic fraction of the culture. By passage 10, isolates grew largely planktonically as indicated by a low percentage of the population growing as an aggregate (percent aggregated). **C.** Whole genome sequencing of experimentally evolved isolates identified mutations at the FMG_0048/0049 locus, including a point mutation in the promoter region (C → T), a nonsynonymous amino acid mutation (T451R), and frameshifts (fs) that result in premature stop codons. Each mutation shown occurred in a distinct evolved isolate, with two isolates having the same mutation in the promoter region.

To learn more about the functions encoded by these genes, we used the SMART domain analysis tool to identify predicted domains within FMG_0048 and FMG_0049. Both proteins have a predicted signal peptide at the N-terminus and a set of Internalin B-like, B-repeat domains (IPR013378), both of which are found in cell surface proteins (37). FMG_0049 featured additional internal repeats and three predicted cell wall anchoring (CWA) domains at the C-terminus (Fig. 4A). Bacterial aggregation is often mediated by large, repetitive cell surface proteins called adhesins, which promote attachment to self or surfaces (38). Based on their size (∼2,500 amino acids), role in aggregation, and predicted domain architecture, we proposed that these genes encode one or two adhesins (Fig. 4A).

**Figure 4.**
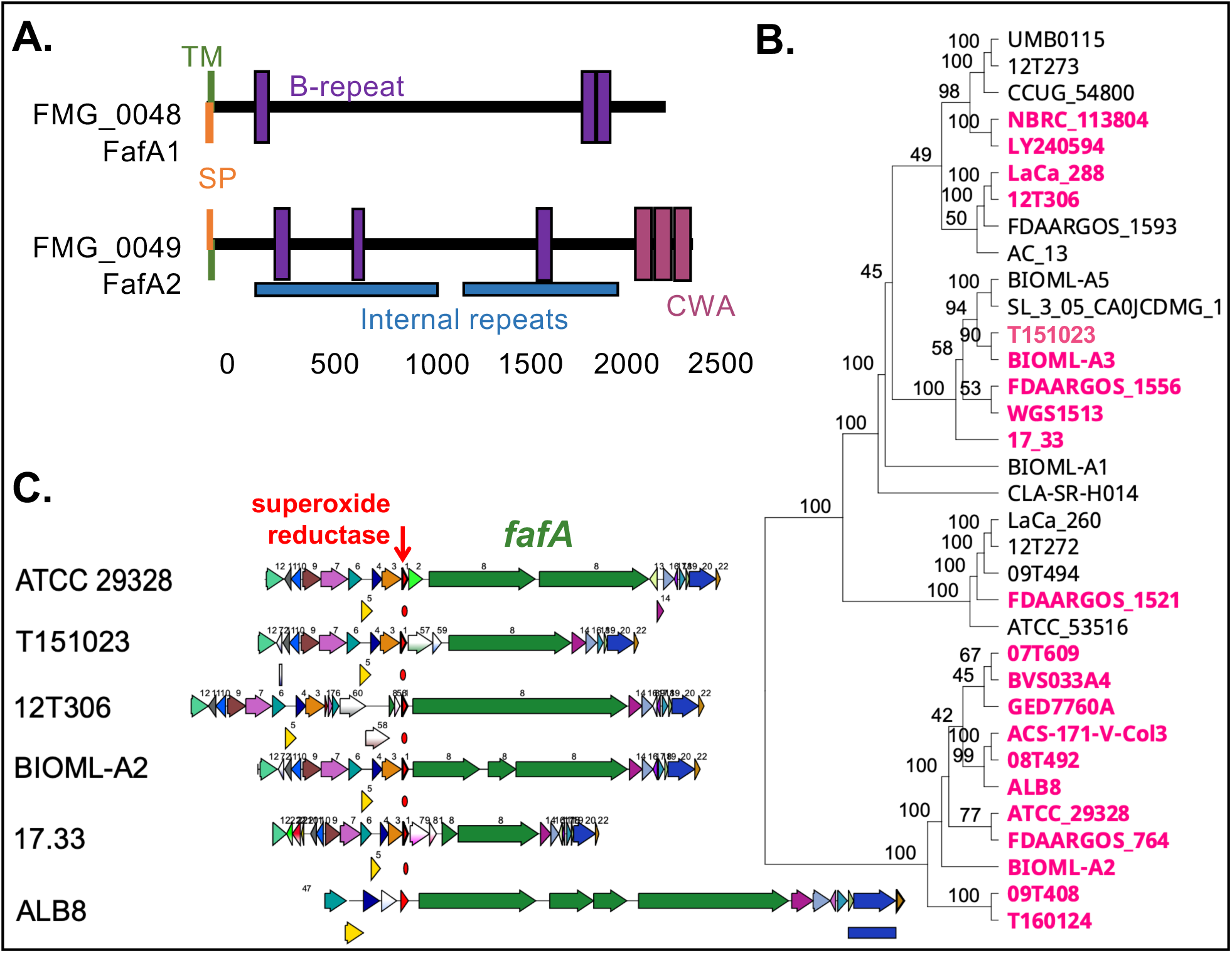
Putative adhesin FafA is widely distributed across the *F. magna* phylogeny but displays variation in gene organization at the conserved locus. **A.** The predicted domain architectures of proteins encoded by FMG_0048 (FafA1) and FMG_0049 (FafA2) include a signal peptide (SP), transmembrane domain (TM), and Internalin B-like B-repeat domains. FafA2 also includes predicted internal repeats and cell wall anchoring domains (CWA). **B.** Phylogenomic tree based on core-genome SNPs of 34 sequenced *Finegoldia* genomes. Strains in pink encode putative FafA homologs. Values show bootstrap percentages for 1000 bootstrap replicates. **C.** Genomic neighborhood surrounding *fafA* in representative *F. magna* strains. Green arrows denote *fafA*, while neighboring arrows represent annotated coding sequences. Conserved synteny is observed among strains carrying the canonical *fafA* locus, which is located downstream of superoxide reductase (red arrow). The *fafA* locus is comprised of one, two, three, or four open reading frames.

Because of their similar domain architecture, we initially considered whether one of these genes resulted from a duplication event. However, when aligned, FMG_0048 and FMG_0049 exhibited low amino acid similarity (35%, Fig. S1A), indicating a recent duplication event did not give rise to these adjacent genes. We next aimed to determine whether homologs were present in other *F. magna* strains. Using tBLASTn, we found that 21 of 34 *F. magna* genomes (62%) had predicted homologs to FMG_0048 (Figs. 4B, S2). Of the 21 identified homologs, 20 occurred at a conserved locus immediately downstream of a gene encoding superoxide reductase (Figs. 4C, S2). For the sole exception, the FMG_0048 homolog occurred within a region containing multiple recombinases and an LtrA reverse transcriptase (Fig. S2), suggesting that this region has been shaped by recombination or mobile genetic element activity. Thus, with rare exception, this locus is generally syntenic across *F. magna*.

We next searched *F. magna* genomes for FMG_0049 homologs. Using FMG_0049 as a query identified the same genomic region as FMG_0048, however this search showed that gene structure varies at this locus across *F. magna* strains. In most strains, FMG_0048 and FMG_0049 both aligned to different regions of a single open reading frame (ORF), while other isolates encoded 3 or 4 ORFs at this location (Fig. 4C, Fig. S2). Of our included strains, only *F. magna* ATCC 29328 encoded 2 ORFs. Among single-ORF loci, the length of the FMG_0048-0049 homolog varied substantially across isolates, ranging from 2,755 to 5,806 amino acids. FMG_0048 aligned to the N-terminal region of single-ORF homologs, whereas FMG_0049 aligned to the C-terminal region (Fig. S1B). Likewise, in multi-ORF loci, FMG_0048 aligned to the first one or two ORFs, whereas FMG_0049 aligned to downstream ORFs (Fig. S1B). Overall, these findings suggest that FMG_0048 and FMG_0049 belong to a conserved but structurally variable locus, consistent with a history of gene fission or fusion.

When we aligned the amino acid sequences of all 21 FafA homologs, we found only specific regions were highly conserved (Fig. S1B). The region corresponding to the N-terminal ∼1,600 amino acids of single-ORF structures, which includes the signal peptide and putative adhesive domain, as well as the C-terminal ∼1,100 amino acids, which includes the cell wall anchoring domain, exhibited higher conservation (30-100% identity) compared to the middle region of the protein which was highly variable (< 30% identity). This pattern further supports the prediction that FMG_0048-FMG_0049 and homologs encode an adhesin, since adhesin structure is often modular, exhibiting conserved N-terminal adhesive and C-terminal cell wall-anchoring domains that are separated by a variable and repetitive stalk region (38, 39).

In prior work, Frick et al. identified FAF (*<u>F</u>. magna* <u>a</u>dhesion factor), a protein implicated in autoaggregation and attachment to the basement membrane protein BM-40 on skin (26). Here, we follow this naming convention by designating this locus *fafA* (*<u>F</u>. magna* <u>a</u>dhesion factor <u>A</u>). Since our analyses indicate that FMG_0048 and FMG_0049 represent regions of a single adhesin locus encoded by one ORF in most *F. magna* isolates, we refer to the ATCC 29328 genes as *fafA1* and *fafA2*, respectively. Having shown that *fafA* is prevalent and broadly distributed across the *F. magna* phylogeny (Fig. 4B), we next aimed to determine whether this adhesin is present in other species. We conducted a BLASTp search against NCBI’s nonredundant bacterial protein database using FMG_0048 and FMG_0049 as queries and found putative homologs in several related obligate anaerobe genera including *Anaerococcus*, *Peptoniphilus*, and *Peptostreptococcus* (Dataset S2). Thus, this adhesin potentially regulates aggregation in other host-associated anaerobes.

### Autoaggregation requires the adhesin, FafA

To verify the role of *fafA1A2* in aggregation, we constructed both single mutants as well as the double mutant by replacing each or both ORFs with *ermB* (Fig. 5A). As noted above, introduction of *ermB* into ATCC 29328 results in loss of its native plasmid. Likewise, whole genome sequencing of *fafA* mutant strains confirmed that they had also lost the native plasmid. Thus, we use the plasmid-free *tuf-ermB* strain of ATCC 29328 constructed above as an additional control in the following experiments, which we hereon refer to as WT^p-^. When grown statically in THB-Tween, WT^p-^ showed robust aggregation (Fig. 5B, C), confirming loss of the native plasmid does not affect aggregation. However, deletion of *fafA1*, *fafA2*, or both resulted in loss of visible aggregation (Fig. 5B) and a marked decrease in percent aggregated cells (Fig. 5C), demonstrating that this locus is required for aggregation under static conditions.

**Figure 5.**
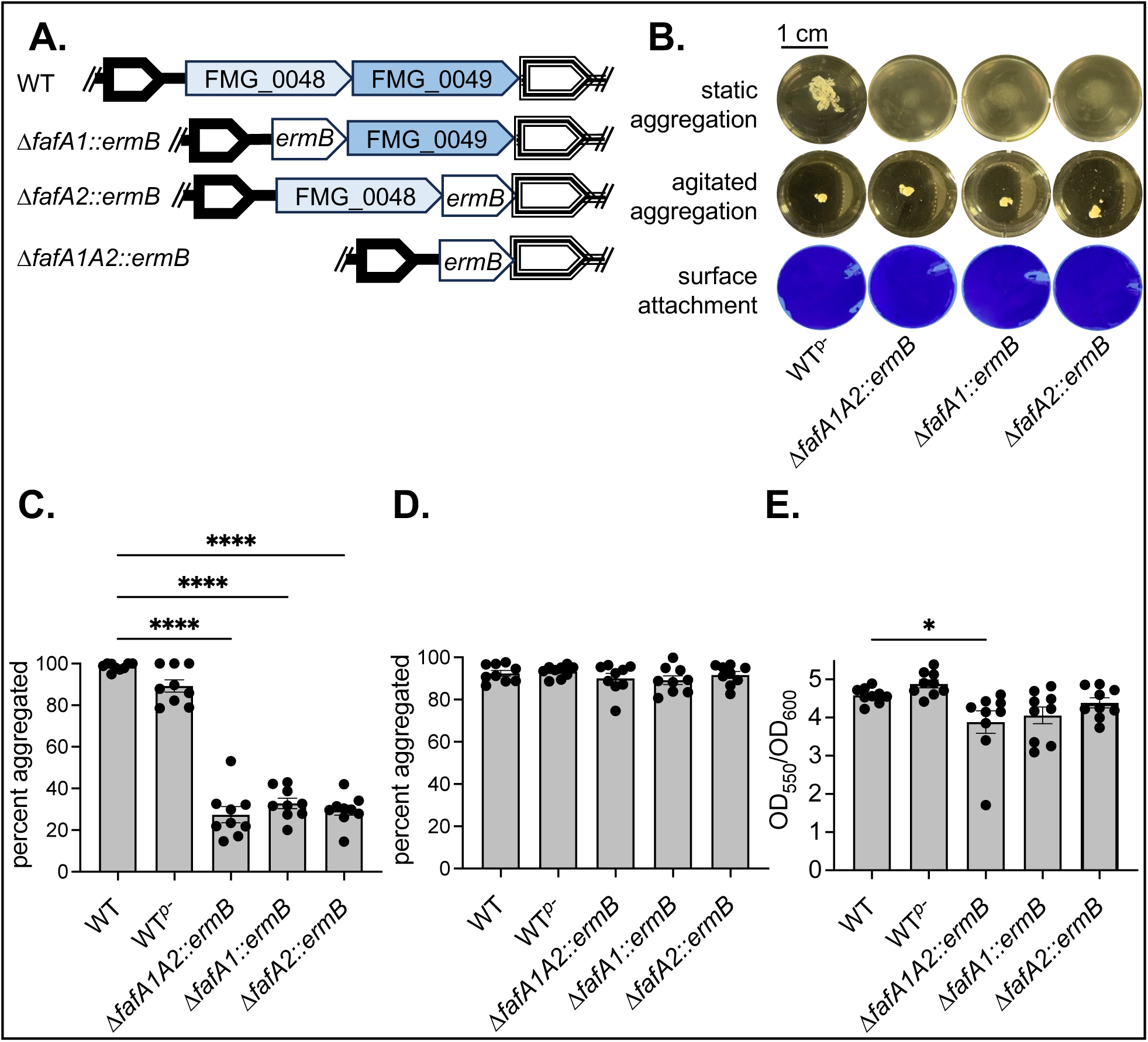
FafA is required for static autoaggregation but not other modes of biofilm formation. **A.** Strains used for biofilm experiments include ATCC 29328 WT (containing a native plasmid) and isogenic mutants that have lost the native plasmid: WT*^p-^* (*tuf-ermB*), Δ*fafA1::ermB, fafA2::ermB, and ΔfafA1A2::ermB*. **B.** Representative images of biofilm phenotypes for mutant strains, including **C.** static aggregation, **D.** agitated aggregation, and **E.** surface attachment, the last of which was quantified as crystal violet stain (OD_550_) normalized to cell growth (OD_600_). Shown are three technical replicates from three independent experiments with the mean ± SEM. Differences in biofilm formation between WT and mutant strains were determined by one-way ANOVA followed by Dunnett’s post-hoc test for multiple comparisons to WT; no stars = not significant, * = *P* < 0.05, ** = *P* < 0.01, *** = *P* < 0.001, and **** = *P* < 0.0001.

In addition to forming aggregates under static growth conditions, we have identified two additional modes of biofilm formation in ATCC 29328 (40). First, when *F. magna* is cultured in THB-Tween with agitation (i.e. continuous shaking at 120 rpm), it forms a more compact aggregate (Fig. 5B). Second, when cultured in a defined medium (DM) in the presence of a glass coverslip, *F. magna* forms a surface-attached biofilm on the glass that is readily visualized with crystal violet staining (Fig. 5B) (40). Interestingly, *fafA1* or *fafA2* are dispensable for these other modes of biofilm formation (Fig. 5B-E). During growth with agitation, WT^p-^ and all deletion strains form aggregates (Fig. 5B, 5D). Similarly, surface attachment was not substantially reduced in the deletion mutants relative to WT (Fig. 5B, 5E). Together, these results demonstrate that *fafA* specifically promotes static autoaggregation in *F. magna* but is not required for other biofilm behaviors.

### FafA-mediated aggregation protects F. magna from antibiotic and oxidative stress

Biofilms are notoriously recalcitrant to killing by antibiotics and host immune defenses (7, 41). To determine whether *fafA*-mediated aggregation protects ATCC 29328 from antibiotic and oxidative stress, we quantified percent survival of WT^p-^ and Δ*fafA1A2*::*ermB* after exposure to metronidazole or H_2_O_2_, the latter of which is generated by immune cells. As expected, the non-aggregating mutant was substantially more sensitive to metronidazole or H_2_O_2_ treatment (Fig 6). At higher concentrations of metronidazole (50 µg/mL) or H_2_O_2_ (0.75 mM), Δ*fafA1A2::ermB* exhibited at least a 3-log reduction in survival compared to WT^p-^. To test whether this protection depended on aggregation, we mechanically disrupted aggregates in WT^p-^ and Δ*fafA1A2::ermB* cultures via vigorous pipetting prior to treatment. Indeed, disruption of WT^p-^ aggregates markedly reduced its survival to metronidazole and H_2_O_2_, often to levels near or below the detection limit. At lower treatment concentrations, physical disruption also further sensitized Δ*fafA1A2::ermB* to these stressors, suggesting cell disruption can increase stress sensitivity in cells already impaired for aggregation. These results demonstrate that *fafA*-dependent aggregation confers protection against both antibiotic and oxidative stress in *F. magna*.

**Figure 6.**
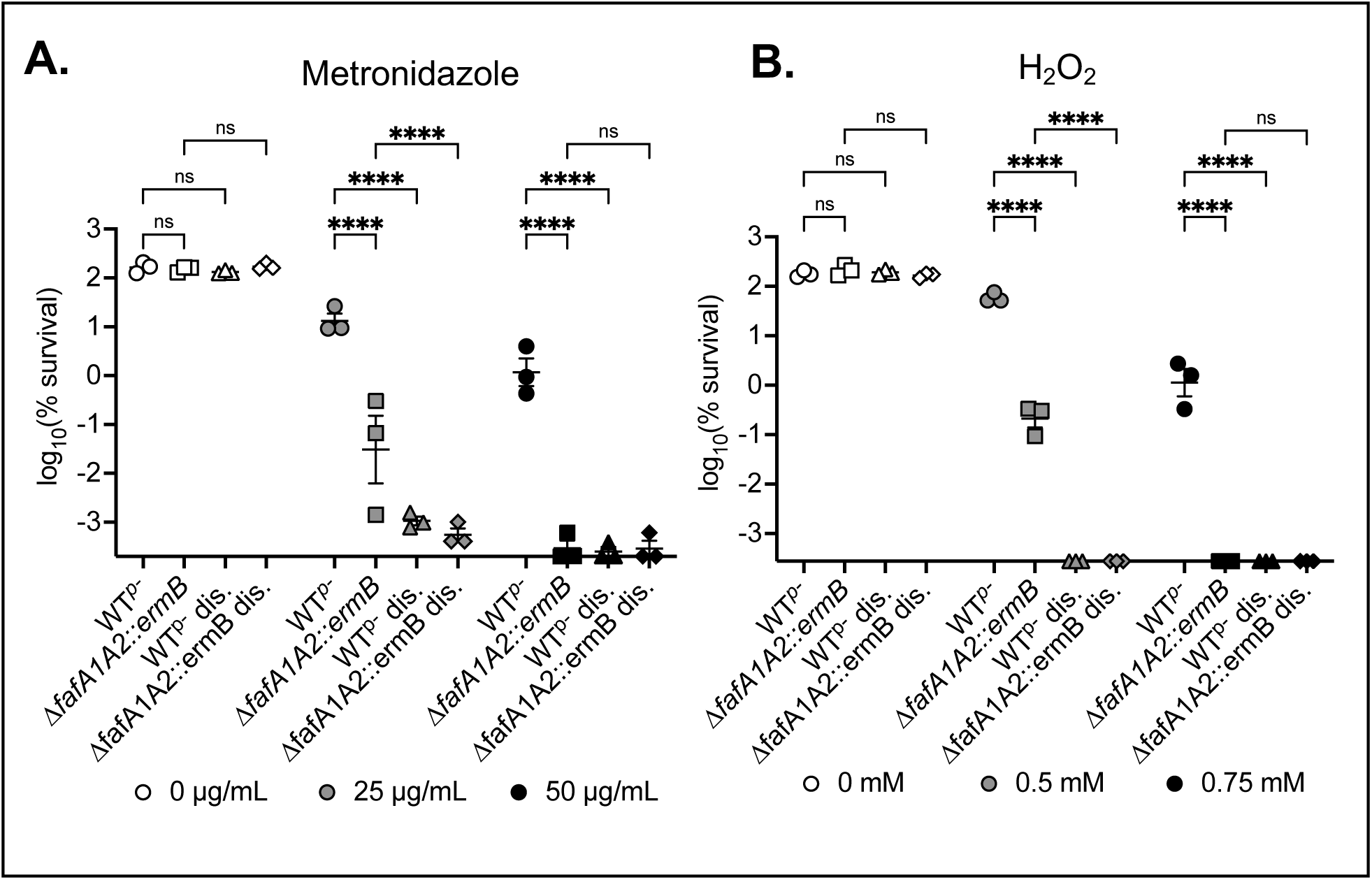
FafA-mediated autoaggregation protects *F. magna* from antibiotic and oxidative stress. The log_10_(%survival) of *F. magna* cultures treated with **A**. 0, 25, or 50 µg/mL metronidazole or **B**. 0, 0.5, or 0.75 mM H_2_O_2_ for 4 hours. Disrupted cultures (dis.) were vigorously pipetted to disrupt aggregates. All data points on the x-axis are below the detection limit of the assay. Data show the means of three replicates, and error bars show SEM. For statistical analyses, data points below the detection limit were conservatively assigned the value of the detection limit. Statistical significance was determined by a two-way ANOVA on log-transformed data followed by Tukey post-hoc test for multiple comparisons; ns = not significant, * = *P* < 0.05, ** = *P* < 0.01, *** = *P* < 0.001, and **** = *P* < 0.0001.

## Discussion

Many clinically relevant bacteria remain genetically intractable, limiting our ability to understand the functions that shape their biology and contributions to human health and disease. In particular, obligate anaerobes dominate the human microbiota and play critical but not fully understood roles in host physiology. Here, we demonstrate a framework for genetic discovery in *Finegoldia magna*, an anaerobic commensal and opportunistic pathogen. Our approach combines complementary techniques: experimental evolution provides an unbiased, forward-genetic approach for identifying genes of interest, and natural competence-mediated mutagenesis enables reverse-genetic studies of gene functions. Overall, this work begins to address a key priority in microbiome research: developing genetic approaches that enable mechanistic studies of host-associated bacteria (42–44).

Many host-associated bacteria encode putative competence genes (45, 46), highlighting natural competence as a potential route for genetic manipulation in these organisms. Other research groups have successfully exploited endogenous natural competence systems to develop genetic tools in understudied bacteria, including Gram-negative anaerobes like *Veillonella* and *Porphyromonas* (47–50), and more recently, in the Gram-positive anaerobe *Parvimonas* (30, 32). Here, we modified and expanded these tools for use in *F. magna*, demonstrating natural competence in several strains, which enabled efficient transformation and targeted mutagenesis. Although *F. magna* genomes encode several putative competence-associated (*com/rec*) genes, the exact mechanism by which *F. magna* takes up and incorporates DNA into its genome is still unknown. Putative *com* and *rec* genes are also present in other closely related anaerobic commensals, including *Anaerococcus* and *Peptoniphilus*, suggesting the full utility of this approach remains to be explored.

Although transformation was successful across multiple *F. magna* strains, its frequency varied considerably among isolates and between liquid- and plate-based incubation protocols. These findings indicate that natural competence in *F. magna* is influenced by both genetic background and environmental factors, as has been observed in other naturally competent organisms (45, 51–59), consistent with competence as a carefully regulated state in bacteria. We also found that transformation frequency increases with the length of homologous flanking regions, which is also known to be an important parameter of competence-based mutagenesis in other bacteria (30, 47, 60, 61).

Beyond its use as a genetic tool, little is known about the role of natural competence in *F. magna* biology. In other bacteria, natural competence facilitates horizontal gene transfer, DNA repair, or nutrient acquisition, although the relative importance of these functions varies by species and environmental context (62–65). Given that *F. magna* occupies dense biofilm communities within polymicrobial chronic infections (2, 25), DNA uptake may provide opportunities for genetic exchange with neighboring organisms. Alternatively, competence may contribute to nutrient acquisition, particularly since nucleosides are one of the few metabolites known to support *F. magna* growth (40). While these ideas remain speculative, future studies should investigate whether competence contributes to genetic exchange or nutrient acquisition in *F. magna* and related anaerobes.

This study also supports the utility of experimental evolution as gene discovery tool for bacteria that lack well-established genetic systems (66–68). Here, phenotype-based selection for the loss of aggregation identified the *fafA* locus, whose role in autoaggregation was validated via targeted mutagenesis. The *fafA* locus varied across isolates with regard to gene organization and protein size, consistent with prior work showing that adhesins often have modular, strain-variable architectures that are altered by domain gain, loss, or rearrangement (69, 70). Although nearly all *fafA* homologs occupied a conserved genomic position, one homolog (strain FDAARGOS_764) was found at a distinct region enriched for genes associated with mobile genetic elements. This observation suggests that the *fafA* locus occasionally undergoes genomic rearrangement or horizontal transfer, although additional analyses would be required to distinguish between these mechanisms. The presence of putative FafA homologs in closely related anaerobes raises the possibility that adhesin genes may be exchanged among host-associated anaerobes, although more rigorous analyses are necessary to distinguish horizontal gene transfer from shared ancestry.

The prevalence of FafA in *F. magna* and relatives suggests that this adhesin makes important contributions to its persistence in the host environment. Indeed, we found that FafA-mediated aggregation enhances *F. magna* tolerance to infection-relevant stresses, including oxidative and antibiotic stress. This finding is in line with a wealth of research in other bacteria, demonstrating that biofilm formation provides protection from a variety of stressors. This protection has been attributed to several mechanisms, including physically shielding cells from antimicrobial compounds and phagocytosis, as well as generating physiological heterogeneity that allows a subset of cells to survive stressful conditions (41). Future studies will examine the mechanism by which FafA-mediated aggregation protects *F. magna*.

We also showed that FafA mediates static autoaggregation, but not other forms of biofilm formation, including agitated aggregation and surface attachment. The observation that FafA mediates a distinct biofilm mode is in line with the field’s increasing appreciation that the term “biofilm” is broad and does not represent a single state (6, 71). The identification of multiple, genetically distinct biofilm lifestyles suggests that biofilm formation is a central feature of *F. magna* biology. These diverse lifestyles may reflect adaptation to the range of host environments occupied by *F. magna*, including the gut, healthy skin, chronic wounds, and the surfaces of implanted medical devices. *F. magna* biofilm formation may be particularly significant in infection contexts, promoting treatment tolerance and impairing immune clearance, which can ultimately delay healing.

Overall, the genetic tools established in this study were used to highlight that *F. magna* encodes multiple modes of biofilm formation, regulated by different genes and environmental cues (40). Future work will continue to expand the genetic toolkit for *F. magna,* including the development of genetic complementation, additional antibiotic resistance markers, and genome-wide approaches such as generation of transposon mutant libraries. Expansion of these tools will enable investigation of new biological questions, such as determining the function of FafA domains and whether they can mediate interactions with other *F. magna* isolates, neighboring microbial species, or host cells. Future studies should also investigate the contribution of FafA to host colonization and infection, testing whether variation among FafA homologs influences biofilm formation and persistence. Similarly, identifying the environmental cues and regulatory pathways that control FafA-dependent aggregation and other biofilm states will illuminate how *F. magna* transitions between different community lifestyles. Ultimately, defining how these distinct biofilm programs shape host colonization, polymicrobial interactions, and infection persistence will provide critical insight into understanding the dual role of *F. magna* as a commensal and opportunistic pathogen.

## Materials and Methods

### Bacterial strains and growth conditions

All strains used in this study are listed in Dataset S3 and include *F. magna* ATCC 29328, 17.33, 09T494, and 07T609, as well as isogenic mutant strains whose construction is described below. ATCC 29328 was isolated from an abdominal wound, 17.33 was isolated from a leg ulcer (strain received from Dr. Ariane Neumann and Dr. Inga-Maria Frick), and 09T494 and 07T609 were isolated from orthopedic implant infections of the hip and foot, respectively (strains received from Lise Hald Schultz and Dr. Holger Brüggemann) (11).

*F. magna* strains were cultured anaerobically at 37°C in either an anaerobic glove box or an anaerobic jar. The anaerobic glove box (Coy Laboratory Products) contains a 95% N_2_ and 5% H_2_ atmosphere. To use the anaerobic jar, cultures were sealed into an AnaeroPack System 2.5 L rectangular jar (Mitsubishi Gas Chemical Co, 50-25); anaerobic jars were either sealed in the anaerobic glove box (maintaining a 95/5% N_2_/H_2_ headspace) or one GasPak EZ Anaerobe Container System Sachet (BD, 260678) was added to the jar prior to sealing to generate an anoxic atmosphere. Liquid cultures of *F. magna* were maintained in Todd Hewitt Broth (Research Products International, T47500) supplemented with 1% Tween 80 (Research Products International, P20390), referred to as THB-Tween. *F. magna* strains were also cultured on THB-Tween blood agar plates made with 4.2% w/v Todd Hewitt Broth powder (RPI), 0.98% w/v NaCl, 0.7% v/v Tween 80, 2.1% w/v agar, and 1% v/v Sheep’s Blood (HemoStat Laboratories), the last of which is added after autoclaving. Where specified, *F. magna* was cultured in a defined medium, described previously (40), supplemented with 60 mM fructose plus 60 mM pyruvate for carbon.

### Cell lysis and purification of F. magna genomic DNA

Genomic DNA was extracted using a modified protocol incorporating mechanical cell lysis prior to purification with the Monarch® Spin gDNA Extraction Kit (New England Biolabs). Briefly, 1 mL of a 10 mL overnight culture (OD_600_ ≈ 0.5) was pelleted by centrifugation and resuspended in 100 μL of 1x TE buffer (10 mM Tris-HCl, 1 mM EDTA). Cells were lysed by bead beating with 0.1-mm glass disruption beads (Research Products International; Cat. No. 9830) for three 30-second cycles at 4,350 rpm. Following mechanical disruption, 100 μL of 2x lysis buffer (40 mM Tris-HCl, pH 8.0; 4 mM EDTA; 2.5% Triton X-100) was added, and samples were incubated at 37°C for 30 min. The samples were then centrifuged to pellet the beads, and the supernatant was transferred to a fresh 1.5-mL microcentrifuge tube. Genomic DNA was purified using the Monarch® Spin gDNA Extraction Kit according to the manufacturer’s instructions, beginning with the addition of 1 μL Proteinase K and 3 μL RNase A to the 200 μL cell lysate.

### Assembly of mutagenesis constructs

All primers are listed in Dataset S1. Primers and constructs were designed using Geneious Prime software (version 2026.0.2, www.geneious.com). Phusion Plus PCR 2X Master Mix (Thermo Fisher Scientific) was used to amplify all DNA fragments for mutagenesis, whereas clone screening was performed via colony PCR using GoTaq 2X master mix (Promega Corporation).

For initial tests of natural competence, an erythromycin resistance cassette (*ermB*) was inserted directly downstream of the elongation factor Tu (*tuf*) gene (FMG_0493). The *ermB* cassette with its native promoter was amplified from pMRMTK-clo10 (provided by Dr. Nathan Crook; Addgene plasmid #203961) using primers with 20-25 bp overhangs with homology to the *F. magna* genome for construct assembly. Upstream and downstream regions (each 2 kb in size) flanking the *tuf* insertion site were each amplified from genomic DNA of the corresponding *F. magna* strain, and PCR products were purified via the Monarch® PCR & DNA Cleanup Kit (New England Biolabs). Next, upstream-*ermB*-downstream constructs were assembled via the NEBuilder HiFi DNA Assembly Cloning Kit (New England Biolabs). Assembled products were PCR-amplified, purified, and used in transformation protocols (see details below).

To generate mutagenesis constructs with different lengths of homologous flanking regions, we amplified *ermB* and the surrounding genomic regions from purified genomic DNA of previously generated ATCC 29328 and 17.33 *tuf-ermB* strains; primer pairs annealing at different distances upstream and downstream of *ermB* were used to vary homologous flank size. To construct ATCC 29328 Δ*fafA1::ermB*, Δ*fafA2::ermB*, and Δ*fafA1fafA2::ermB*, we assembled a DNA construct of *ermB* flanked by 2 kb fragments homologous to the upstream and downstream regions of *fafA1*, *fafA2*, and *fafA1/A2*, respectively. As described above, assembled fragments were PCR amplified, purified, and used for transformation via the protocol described below.

### Natural competence-based transformation in F. magna isolates

For initial studies of transformation frequency, including those testing the effect of homologous flank size and DNA concentration, we transformed cells using the spot plate incubation method, modified from Higashi et al (30). In this method, overnight cultures were resuspended in fresh THB-Tween medium to OD_600_ of 0.4, and 40 µl of the cell resuspension was mixed with 1 µg of purified PCR product (or the molar equivalent for different flank sizes) resuspended in 5 – 10 µl of sterile water (final volume of 45 – 50 µl). The cell-DNA suspension was then spotted onto a THB-Tween blood agar plate, which was incubated at 37°C anaerobically overnight. Following incubation, biomass from the cell spot was collected with a sterile swab, resuspended in 500 µl PBS, and serially diluted in PBS before plating onto THB-Tween blood agar and THB-Tween blood agar with 15 µg/mL erythromycin. Transformation frequency was calculated as the CFU on erythromycin plates divided by the CFU on the antibiotic-free plates. Several transformants were screened via colony PCR to confirm they contained the *ermB* insert. Finally, whole genome sequencing (Plasmidsaurus) was performed on one transformant from each strain, with custom analysis and annotation.

As specified, some transformation experiments were conducted via a liquid incubation method. As above, overnight cultures of *F. magna* were resuspended in fresh THB-Tween medium to OD_600_ of 0.4. Next, 100 µl of cell suspension was mixed with 10 µl of purified PCR product (ranging from 0.1 - 2 µg), and the cell-DNA suspension was incubated in a 1.5 mL Eppendorf tube at 37°C anaerobically overnight. Cells were serially diluted in PBS and plated onto THB-Tween blood agar plates with or without 15 µg/mL erythromycin to determine transformation frequency, as described above.

### Growth and quantification of static or agitated aggregates

To grow *F. magna* as an aggregate under static conditions, overnight cultures of ATCC 29328 were diluted in fresh THB-Tween medium to OD_600_ of 0.005 as 2 mL cultures in a 12-well plate. Cultures were incubated at 37°C for 24 hours to allow for growth, after which cultures were gently swirled for 15 seconds to promote autoaggregation and dislodge any planktonic cells. Culture plates were then incubated statically for 15 minutes to allow aggregates to settle.

To grow agitated aggregates, overnight ATCC 29328 cultures were diluted in fresh THB-Tween medium to OD_600_ of 0.005 as 2 mL cultures in a 12-well plate. Cultures were grown at 37°C in an anaerobic jar (95/5% N_2_/H_2_ atmosphere) that was placed on a shaking incubator (37°C, 120 rpm).

To quantify the extent of aggregation, the percent of culture biomass that was aggregated was calculated. Percent aggregated was calculated by subtracting the OD_600_ of planktonic cells (OD_planktonic_) from the OD_600_ of the entire culture (OD_total_), thereby determining OD_aggregated_, which is ultimately divided by OD_total_ and multiplied by 100 to yield percent aggregated. To measure OD_planktonic_, 1 mL of the undisturbed culture (avoiding the aggregate) was transferred into a cuvette for OD_600_ measurement. Planktonic cells were then transferred from the cuvette back into the original culture well, and the entire culture was pipetted vigorously to disrupt aggregates. To measure the OD_total_, 1 mL of the disrupted culture was transferred to a cuvette and the OD_600_ was measured.

### Growth and quantification of surface attached biofilms

To grow surface attached biofilms, ATCC 29328 was grown in defined medium (DM) in the presence of glass coverslips, as previously (40). Briefly, overnight cultures were pelleted and washed twice with PBS to remove residual THB-Tween medium. Washed cells were inoculated into fresh DM at OD_600_ of 0.01 and incubated as 2 mL cultures in a 12-well plate, where each well contained a sterile glass coverslip (VWR, #16004-300). Culture plates were placed in an anaerobic jar with one sachet (BD, 260678) and incubated statically for 72 hours at 37°C to allow for biofilm formation on coverslips. To quantify surface-attached biofilms, glass coverslips were removed from wells using sterile tweezers and non-adhered cells were removed by gently swirling coverslips three times in water. Coverslips were stained in a 0.1% crystal violet solution for 1 minute, and subsequently washed four times in water to remove unbound stain. Stained coverslips were placed in wells of a 6-well plate and incubated with 1 mL of 30% acetic acid on a gentle plate rotator for 10 minutes to solubilize the crystal violet stain. The OD_550_ of each sample was measured to quantify crystal violet stain, using 30% acetic acid as a blank. Crystal violet measurements (OD_550_) were normalized to bacterial growth (OD_600_), the latter of which was determined using one parallel unstained culture per condition wherein cultures were vigorously pipetted to remove surface attached cells and quantify total growth (OD_600_). Thus, surface attachment is reported as OD_550_/OD_600_.

### Experimental evolution to identify fafA

To select for mutants that were incapable of forming static aggregates, ATCC 29328 cultures were grown statically overnight, after which only non-aggregating cells (i.e. from the planktonic portion of cultures) were serially passaged into fresh THB-Tween medium and incubated for growth. For each of 10 experimental lineages, 20-200 uL of planktonic cell populations was passaged every 24-72 hours, with cultures maintained under static conditions. After 10 passages, all experimental lineages grew largely planktonically, and isolates from each lineage were streak purified. Whole genome sequencing of evolved isolates from several lineages was performed by Plasmidsaurus. Mutations were identified in the evolved isolates by comparing their genome to the parent strain via breseq, with default settings (Table S1) (72).

### FafA domain architecture, homolog identification, and phylogenetic mapping

To better understand FafA function, protein domain architecture was predicted using SMART (Simple Modular Architecture Research Tool) (73). Amino acid sequences of FMG_0048 and FMG_0049 were analyzed with searches against the SMART and Pfam databases, including predictions of signal peptides and internal repeats. Predicted domains shown in Fig. 4A include domains above the default significance threshold.

Putative homologs of FMG_0048 and FMG_0049 were identified across *F. magna* strains using tBLASTn searches against a custom BLAST database containing 34 *F. magna* RefSeq genomes (Dataset S3). Searches were performed in Geneious Prime. Using FMG_0049 as query required removal of the cell wall anchoring domain sequence (281 bp) from the C-terminus, as this region was conserved across variety of cell surface proteins. Only BLAST hits with an E-value below 1 x 10^-10^ were further considered as potential homologs. To identify putative FMG_0048 and FMG_0049 homologs across other bacteria (i.e. outside of *Finegoldia*), BLASTp was used against the NCBI bacterial nonredundant protein database with strict cutoffs of ≥60% query coverage and ≥60% sequence identity (Dataset S2).

Having identified putative *F. magna* FafA homologs via tBLASTn, we next used synteny analysis as additional evidence of FafA homology across *F. magna*. Genomic neighborhoods surrounding each putative homolog were visualized in Geneious Prime by extending BLAST hits to include neighboring annotated genes. Conserved synteny was assessed manually by comparing local gene order across isolates and additionally visualized using the BV-BRC compare genome regions tool (Fig. S2) (74). Putative FafA homologs that were located 0-2 ORFs downstream of superoxide reductase were considered to have conserved synteny. For the one putative FafA homolog found at a different genomic location (strain FDAARGOS_764), neighboring genes were manually inspected using RefSeq annotations to identify features associated with genome mobility, such as recombinases and reverse transcriptases.

To compare FafA, FafA1, or FafA2 sequences, pairwise amino acid alignments were generated using MAFFT implemented in Geneious Prime. Global alignments were performed using the BLOSUM62 substitution matrix, a gap opening penalty of 1.0, and an offset value of 0. Amino acid similarity and identity values were calculated from the resulting alignments in Geneious Prime. For multi-ORF loci, concatenated amino acid sequences were used (Fig. S1).

To map the presence/absence of FafA homologs onto the *F. magna* lineage, a whole-genome phylogeny was constructed from 34 *F. magna* RefSeq genomes using Parsnp (version 2.1.4) with *F. magna* ATCC 29328 as the reference genome and the option -c to force inclusion of all 34 genomes (75). The resulting core genome alignment was fed into RAxML-NG (76) to create a maximum likelihood tree with 1000 bootstraps. The resulting tree was visualized in Geneious Prime.

### Stress tolerance assays

To assess the impact of aggregation on antibiotic and oxidative stress tolerance, overnight cultures of WT^p-^ and Δ*fafA1A2::ermB* were diluted into fresh THB-Tween media at OD_600_ of 0.05 as 2 mL cultures in 12-well plates. Cultures were incubated statically at 37°C for 4 hours to allow cells to grow as either aggregates (WT^p-^) or planktonically (Δ*fafA1A2::ermB*). After 4 hours, parallel cultures of each strain were mechanically disrupted (via pipetting) and dilution plated on THB-Tween blood agar to determine initial viable cell counts. Remaining replicate cultures were incubated statically with or without stress at 37°C for 4 hours. For antibiotic treatment, metronidazole (Thermo Scientific #210340050) was added to cultures at a final concentration of 0 (control), 25, or 50 µg/mL. For oxidative stress treatment, H_2_O_2_ (30% solution, Macron Fine Chemicals #5240-05) was added to cultures to achieve final concentrations of 0 (control), 0.5, or 0.75 mM. After 4 hours of stress exposure, cultures were disrupted via pipetting and cells were plated to determine final viable cell counts. Percent survival was quantified by dividing final viable cell counts by the average of initial viable cell counts and multiplying by 100, within each strain.

To test the role of aggregation in stress tolerance, parallel cultures of both WT^p-^ and Δ*fafA1A2::ermB* were vigorously pipetted to disrupt any aggregates. Following disruption, cultures were incubated statically for 20 minutes to allow cells to settle, after which metronidazole or H_2_O_2_ was added to cultures. As above, disrupted cultures were incubated with the stress for 4 hours before plating to quantify percent survival.

## Supporting information

Dataset S1 Primers Table

Dataset S2 fafA1_blastP_nr_bacteria

Dataset S3 strains list

## Acknowledgements

This work was supported by grants to MAS from National Institutes of Health (R35GM155575) and a New Investigator Grant from the Medical Research Foundation of Oregon. Research reported in this publication was supported by the National Institute of General Medical Sciences of the National Institutes of Health under award number T32GM149387. AAC was supported by National Institutes of Health grant R25GM152322. This work was also supported by National Institutes of Health grant K99DK137017 to TJS. We thank Dr. Caitlin Kowalski and Kendal Tinney for assistance developing the *F. magna* lysis protocol. We thank Peter Newstein for help with Inkscape and coding. We thank Connor Siggins for help with figure illustration. We thank Dr. Nathan Crook for generously gifting pMRMTK-clo10. We thank PeaceHealth Sacred Heart Medical Center RiverBend and Dr. Dustin McKague for providing the human wound aspirate from which *F. magna* strain AC_13 was isolated. We thank Dr. Ariane Neumann, Dr. Inga-Maria Frick, Lise Hald Schultz, and Dr. Holger Brüggemann for generously sharing *F. magna* strains that were used in this study.

## Notes

### Competing Interest Statement

The authors have declared no competing interest.

### Summary of Updates

Supplemental materials were added, which were not included in the initial submission. The author list was amended and several references were added to the manuscript text.

