## Supplementary Tables and Figures for "New genetic tools in *Finegoldia magna* identify a conserved adhesin required for the formation of stress-tolerant aggregates"

**Table S1.** Mutations identified via breseq analysis in sequenced isolates purified from passage 10 of experimental evolution selecting for non-aggregating cells. Genome position, locus tag, and predicted product are based on the NCBI annotated genome of *F. magna* ATCC 29328 (AP008971). Bolded are mutations associated with the *fafA* locus.

| Isolate | Genome Position | Mutation Class | Mutation Description | Locus Tag | Product |
|---|---|---|---|---|---|
| **10B1** | | | | | |
| | **52,935** | **Point** | **intergenic (+318/-99) C→T** | **FMG_0047 → / → FMG_0048** | **conserved hypothetical protein/ conserved hypothetical protein** |
| | 831,376 | Non-synonymous point | E305K (GAA→AAA) | FMG_0765 | exopolyphosphatase family proteins |
| **10C1** | | | | | |
| | **52,935** | **Point** | **intergenic (+318/-99) C→T** | **FMG_0047 → / → FMG_0048** | **conserved hypothetical protein/ conserved hypothetical protein** |
| | 128,238 | Non-synonymous point | S728I (AGC→ATC) | FMG_P0138 | hypothetical protein |
| **10E1** | | | | | |
| | **63,423** | **Insertion** | **P1028-T1029 +CC (CCTACA → CCCCTACA)** | **FMG_0049** | **putative N-acetylmuramoyl-L-alanine amidase** |
| **10H1** | | | | | |
| | **53,369** | **Single base deletion** | **E111-K112-A113 (A)$_{5\rightarrow4}$ (GAAAAAGCT→ GAAAAGCT)** | **FMG_0048** | **conserved hypothetical protein** |
| | 1,060,513 | Non-synonymous point | R91R (CGT→CGG) | FMG_0978 | multidrug ABC transporter |
| | 1,501,860 | Non-synonymous point | A17S (GCA→TCA) | FMG_1360 | Phosphoribosylaminoimidazolecarboxamide formyltransferase |
| **10K1** | | | | | |
| | **54,385** | **Non-synonymous point** | **T451R (ACG→AGG)** | **FMG_0048** | **conserved hypothetical protein** |
| | 158,761 | Non-synonymous point | T276I (ACA→ATA) | FMG_0136 | adenylosuccinate synthetase |
| | 1,295,510 | Non-synonymous point | N70I (AAT→ATT) | FMG_1188 | conserved hypothetical protein |
| | 1,501,860 | Non-synonymous point | A17S (GCA→TCA) | FMG_1360 | Phosphoribosylaminoimidazolecarboxamide formyltransferase |

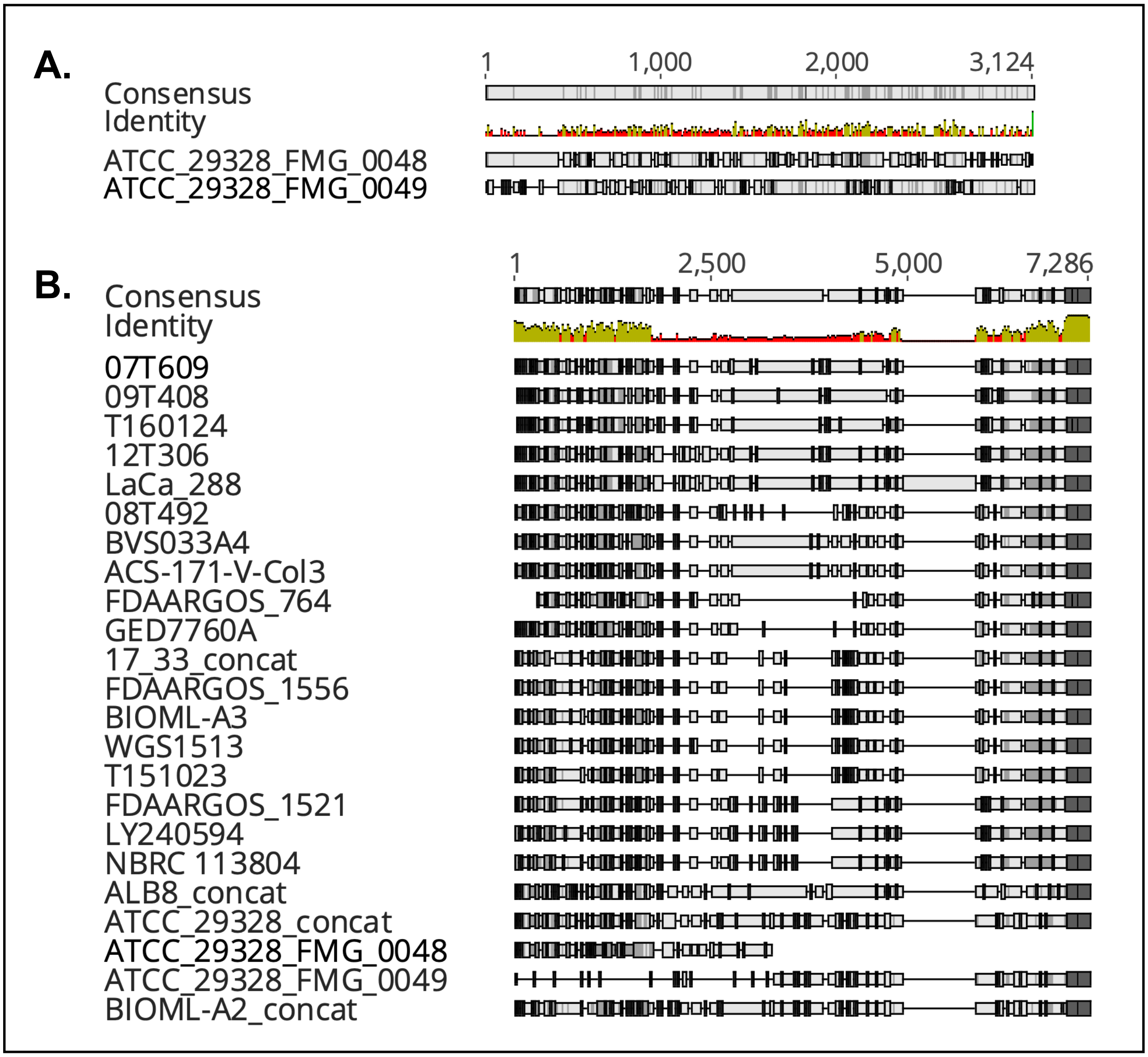


**Figure S1.** MAFFT amino acid alignments of **A.** FMG_0048 and FMG_0049 of ATCC 29328 and **B.** all 21 FafA homologs. For homologs with more than one ORF (17.33, ALB8, BIOML-A2, and ATCC 29328) concatenated amino acid sequences were used. Individual amino acid sequences of FMG_0048 and FMG_0049 from ATCC 29328 were also included to demonstrate suggested history of gene fission/fusion. Mean pairwise identity over all pairs in the column represented by green (100% identity), yellow (30-100% identity), or red (<30% identity).

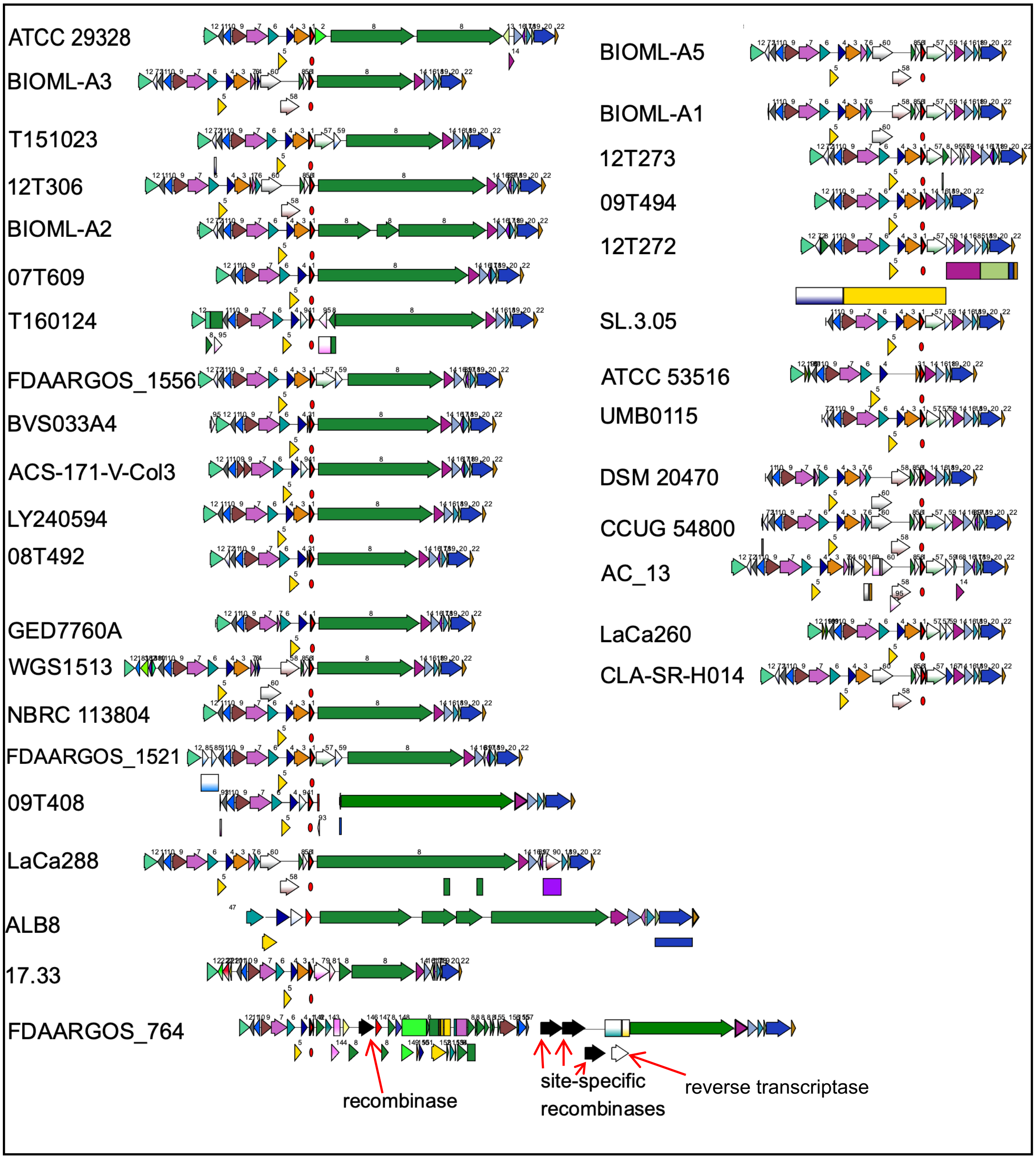


**Figure S2**. Genomic organization of *fafA* locus across *F. magna* strains. *fafA* homologs are depicted as green arrows (labeled "8") and are either absent (right side) or present (left side) as one, two, three, or four open reading frames. All *fafA* homologs are downstream of superoxide reductase gene in red (labeled "1") with red circle, except in FDAARGOS_764.
